# Beyond Lux: Methods for Species and Photoreceptor-Specific Quantification of Ambient Light for Mammals

**DOI:** 10.1101/2023.08.25.554794

**Authors:** Richard J McDowell, Altug Didikoglu, Tom Woelders, Mazie J Gatt, Roelof A Hut, Timothy M Brown, Robert J Lucas

## Abstract

**Background:** Light is a key environmental regulator of physiology and behaviour. Mistimed or insufficient light disrupts circadian rhythms and is associated with impaired health and well-being across mammals. Appropriate lighting is therefore crucial for indoor housed mammals. The most commonly used measurement for lighting is lux. However, this employs a spectral weighting function based on human perceived brightness and is not suitable for ‘non-visual’ effects of light or use across species. In humans, a photoreceptor-specific (α-opic) metrology system has been proposed as a more appropriate way of measuring light.

**Results:** Here we establish technology to allow this α-opic measurement approach to be readily extended to any mammalian species, accounting for differences in photoreceptor types, photopigment spectral sensitivities, and eye anatomy. Since measuring photopigment spectral sensitivity can be hard to derive for novel animals and photoreceptors, we developed a high-throughput, easy-to-use, method to derive spectral sensitivities for recombinantly expressed melanopsins and use it to establish the spectral sensitivity of melanopsin from 12 non-human mammals. We further address the need for simple measurement strategies for species-specific α-opic measures by developing an accessible online toolbox for calculating these units and validating an open hardware, low-cost, multichannel light sensor for ‘point and click’ measurement. We finally demonstrate that species-specific α-opic measurements are superior to photopic lux as predictors of physiological responses to light in mice and allow ecologically relevant comparisons of photosensitivity between species.

**Conclusion:** Our study demonstrates that measuring light more accurately using species-specific α-opic units is superior to the existing unit of photopic lux and holds the promise of improvements to the health and welfare of animals, scientific research reproducibility, agricultural productivity, and energy usage.

## 1. Background

Light is a crucial environmental factor that allows vision and plays a fundamental role in regulating physiological and behavioural processes(1). Light impacts animal biology by setting the phase of circadian rhythms and via direct effects on numerous aspects of behavioural and physiological state(2, 3). Understanding the effects of light on mammalian biology is therefore an important topic of research in its own right(4–7), while ensuring appropriate lighting is an important element of husbandry and a determinant of reproducible outcomes for common experimental paradigms(8–12). The most accessible method of measuring ambient light is to use a lux meter. These are widely obtainable and easy to use. Accordingly, light is commonly quantified in lux in animal research and husbandry(13–15). However, that approach is prone to error, as lux meters employ a light sensor and spectral filtering that match the spectral sensitivity of human flicker photometry (an assay of perceived brightness under cone-favouring conditions)(16). Given this narrow definition of spectral sensitivity, it is unsurprising that lights differing in spectral composition can have quite different impacts on animal biology even if matched for lux(17–19).

The challenge of achieving a wider quantification of ambient light than provided by a lux meter was recently addressed with publication of a new SI-compliant measurement system for light(20, 21). This new metrology aims to quantify light not in relation to its ability to elicit any particular biological response (e.g. perceived brightness in humans) but rather in terms of its effective intensity for each of the retinal photoreceptors responsible for detecting light.

In the case of humans, this allows quantification of 5 different ‘α-opic irradiances’ (rod-opic, melanopic, S-cone-opic, M-cone-opic, and L-cone-opic), corresponding to the effective irradiance for each of the 5 retinal photoreceptors in our own species(20). The definition of α-opic irradiance lends itself to adaption to other species. α-opic irradiance is calculated by weighting energy across the spectrum according to the wavelength sensitivity of the target photoreceptor. It follows that ‘α-opic irradiance’ may be calculated for any photoreceptor of known spectral sensitivity(20). Moreover, the α-opic standard encompasses an additional concept, that of ‘equivalent daylight illumination’ (EDI), which aides cross-species comparisons of light intensity. EDI is the quantity of daylight (in lux) required to produce the corresponding α-opic irradiance(22). A worked example illustrates how this facilitates ethologically relevant comparisons across species: say an experiment in mouse reveals impacts on learning at melanopic irradiance > 1 W/m^2^. Expressing this in terms of melanopic EDI (>500 lux) introduces a relation to an environmental condition (an amount of daylight) which could be experienced by any species. If another study shows that in, say, horses effects on learning are observable only at >1000 lux melanopic EDI, then one could conclude that mice are twice as sensitive as horses to natural light and make precise predictions of the conditions under which light impacts learning in each species.

Although conceptually straightforward, there are currently several practical barriers to widespread adoption of the α-opic metrology across species. The first is incomplete knowledge of photoreceptor spectral sensitivity in some species. To calculate α-opic irradiance we need first to know what wavelength-dependent filter to apply, and this is defined by the wavelength sensitivity of that particular photoreceptor in that species. The second barrier is the absence of simple measurement methods (equivalent of a lux meter) returning α-opic quantities. Here we address these problems. We describe an accessible method for determining unknown photoreceptor wavelength sensitivities and combine this with a review of published data to provide spectral sensitivity functions for calculating α-opic EDIs in major domestic mammal species. We provide an online resource for performing these calculations according to the methodology specified in CIE S026 and show that an open hardware multichannel light sensor can be used to quantify common light sources in α-opic metrics with reasonable accuracy.

## 2. Results

### 2.1. Defining photoreceptor spectral sensitivity

The α-opic metrology weights light energy across the spectrum according to the spectral sensitivity of each class of rod, cone or melanopsin photoreceptor. It follows that defining the spectral sensitivity of these photoreceptors in the target species is the critical step in adapting this approach to use across species. The photoreceptor spectral sensitivity itself is determined by two processes: the fundamental spectral efficiency of the photopigment responsible for light absorption; and the cumulative spectral filtering property of all elements upstream of that photopigment in the light path (pre-receptoral filtering).

A literature review reveals spectral sensitivity information is already available for many photoreceptors across mammalian species (**Supplementary table 1**). The primary exception is melanopsin, whose spectral sensitivity has so far been described in only a few instances. As melanopsin is hard to purify in sufficient quantities for absorbance spectroscopy, we adapted a heterologous action spectroscopy method(23) to determine melanopsin spectral sensitivity for a wider array of species. In brief, HEK293T cells were transfected with an expression vector containing the cDNA sequence of the target melanopsin, presented with an appropriate isoform of retinal as chromophore, and the melanopsin-dependent light response was quantified using a luminescent reporter (**Figure 1A**). Luminescent response amplitude in this assay is dependent on the intensity and spectral composition of the stimulus, providing an opportunity to describe photopigment sensitivity as a function of wavelength(23). Here we exposed cells to 6 spectrally distinct stimuli over a range of irradiances (**Figure 1B**) and used a boot-strap modelling approach to determine the λ_max_ of a putative opsin photopigment which could best predict response amplitude across these stimuli (see methods). When applied to human melanopsin this method returned a λ_max_ estimate of 481 ±1.1nm, similar to published estimates for this species(23, 24) (**Figure 1D**). As a sense check, we confirmed the ability of a pigment with these characteristics to predict the luminescence responses, by plotting response amplitude as a function of effective irradiance for this pigment (weighting irradiance across the spectrum according to pigment sensitivity; **Figure 1E,F**). Having confirmed the suitability of this approach for human melanopsin, we tested it with 3 further melanopsins of known λ_max_ (mouse, crab-eating macaque, and brown rat). In each case our process returned λ_max_ ∼480nm (mouse = 480±1.1nm; macaque = 483±1.2nm; rat = 481±1.1nm) (**Table 1; Supplementary table 2**), closely matching published spectral sensitivity estimates for melanopsin in these species(25–27).

**Figure 1:**
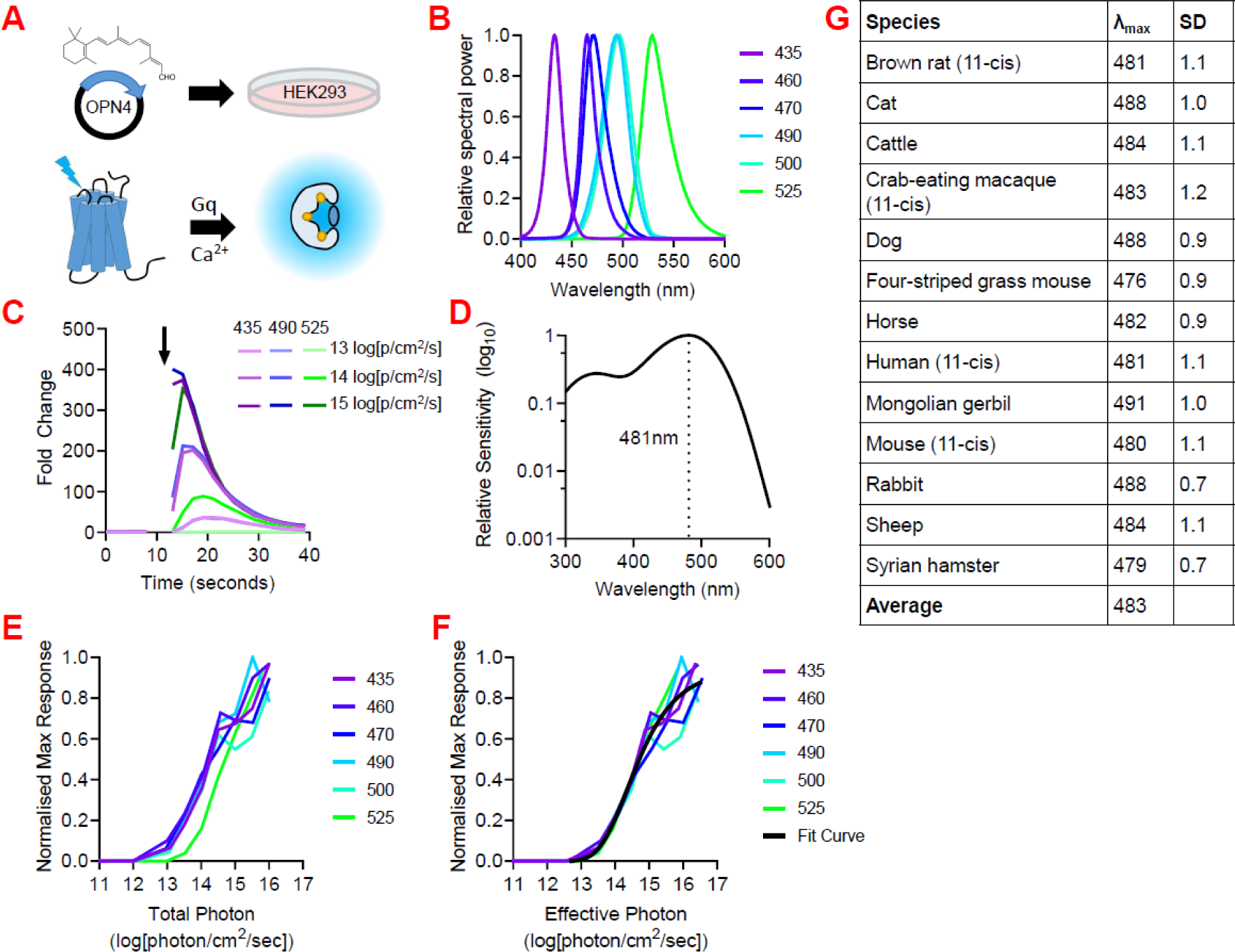
Mammalian melanopsin spectral sensitivities. (**A**) Schematic of action spectra generation. HEK293 cells are incubated with 11-*cis* or 9-*cis* retinal and transfected with plasmid DNA containing melanopsin from the species of interest. Light stimulation drives an increase in intracellular Ca^2+^ via Gq pathway activation, which causes bioluminescence from the Ca^2+^ indicator mtAequorin. Bioluminescence is detected by a plate reader. (**B**) Spectra of stimulating lights used to generate action spectra. (**C**) Example time course showing changes in hOPN4-mediated increases in Ca^2+^ bioluminescence under different stimulating light spectra and intensities. (**D**) Example Govardovskii template for hOPN4 based on predicted λ_max_ 481nm. (**E**) Example irradiance response curves (IRCs) for hOPN4 plotted against uncorrected total photon light intensity. (**F**) Example irradiance response curves (IRCs) for hOPN4 plotted against corrected effective photon light intensity weighted for a photopigment with λ_max_ 481nm. (**G**) Predicted λ_max_ of mammalian melanopsins. Data collected with 9-*cis* retinal and subsequently scaled to λ_max_ for 11-*cis* retinal, unless labelled with ‘(11-*cis*)’, indicating data was collected with 11-*cis* retinal.

**Table 1:** List of mammal photopigment spectral sensitivities.

| Species | Species<br>(latin name) | S-cone-<br>opsin<br>$\lambda_{\max}$ | Melano-<br>psin<br>$\lambda_{\max}$ | Rhodop-<br>sin $\lambda_{\max}$ | M-cone-<br>opsin<br>$\lambda_{\max}$ | L-cone-<br>opsin<br>$\lambda_{\max}$ |
| --- | --- | --- | --- | --- | --- | --- |
| Mouse | Mus musculus | 358 | 480 | 498 | 508 | --- |
| Four-striped<br>grass mouse | Rhabdomys pumilio | 360 | 476 | 493 | 501 | --- |
| Brown rat | Rattus norvegicus | 358 | 481 | 498 | 509 | --- |
| Syrian<br>hamster | Mesocricetus<br>auratus | --- | 479 | 502 | 504 | --- |
| Mongolian<br>gerbil | Meriones<br>unguiculatus | 360 | 491 | 502 | 490 | --- |
| Cattle | Bos taurus | 435 | 484 | 500 | 553* | --- |
| Sheep | Ovis aries | 440* | 484 | 500<sup>#</sup> | 549* | --- |
| Horse | Equus ferus caballus | 428 | 482 | 499 | 545 | --- |
| Cat | Felis catus | 450 | 488 | 501 | 553 | --- |
| Dog | Canis lupus familiaris | 428* | 488 | 506 | 554* | --- |
| Rabbit | Oryctolagus<br>cuniculus | 421* | 488 | 502 | 509 | --- |
| Crab-eating<br>macaque | Macaca fascicularis | 415 | 483 | 500 | 535 | 567 |

<sup>#</sup> Accepted Rod 500nm
\* Adjusted for lens transmission because the raw data come from *in vivo* ERG
References and measurement method in **Supplementary table 1**
Human photopigment spectral sensitivities are based on the CIE metrics for non-visual human light exposure(21): S-cone-opic 447nm, melanopic 488nm, rhodopic 504nm, M-cone-opic 540nm, L-cone-opic 565nm.

Opsin photopigments can employ different *cis* isoforms of retinal as chromophore. As the identity of the isoform alters pigment λ_max_, and as retinoids may be present in culture medium, we finally wished to confirm that the outcome of our assay was determined by the retinaldehyde added to the medium. To this end, we repeated the assay for our 4 test species but adding 9-rather than 11-*cis* retinaldehyde to the culture medium. In other opsins, 9-*cis* retinaldehyde causes a blue shift in spectral sensitivity(28), and this was also the case for melanopsin with a mean±SD short wavelength shift of 16±2.5nm (range = 14 to 20nm) across human, mouse, macaque and rat melanopsins (**Supplementary table 2**).

Having fully validated our approach, we turned to using it to define melanopsin spectral sensitivity for 9 additional domestic mammalian species. To ensure consistency with future work which may make use of commercially available 9-*cis* (in place of harder to obtain 11-*cis*) retinaldehyde, we used the 9-*cis* chromophore for these experiments and applied a 16nm correction. In all cases, the predicted λ_max_ for the 11-*cis* retinaldehyde photopigment was close to 480nm (mean = 483nm; range 476-491nm; **Figure 1G**).

We next turned to the problem of how to account for the contribution of pre-receptoral filtering on photoreceptor spectral sensitivity *in vivo*. In principle this can only be achieved by measuring spectral transmittance of every element upstream of the photoreceptor in the light path, or by describing the wavelength sensitivity of the target photoreceptor *in vivo*. The latter approach has been used in a limited number of species for melanopsin but is not readily applicable to new species. Turning to the former, we identified published reports of spectral transmission for cornea, lens, and vitreous humor for seven mammalian species (**Supplementary figure 1**). Each component of ocular media in all species had good transmission across longer wavelengths. The extent of filtering at shorter wavelengths was species-dependent and predominantly determined by the lens(18, 29). As lens transmission is described for many mammalian species (**Supplementary data 1**) we wondered whether accounting for this parameter alone may adequately predict *in vivo* spectral sensitivity. To this end we identified species in which there was information available for lens transmittance; the absorbance spectrum of purified photopigment; and *in vivo* photoreceptor spectral sensitivity. In these cases, we calculated hypothetical *in vivo* photoreceptor λ_max_ as the product of photopigment *in vitro* spectral sensitivity and lens transmission. In all cases this estimated *in vivo* λ_max_ was similar to the experimentally measured value (**Supplementary table 1**) providing confidence that lens transmittance alone provides a reasonable approximation of pre-receptoral filtering in mammals. The literature contains information about lens transmission for at least 56 mammalian species, including humans (**Supplementary data 1**). The wavelength at 50% transmission (in which higher values represent low UV transmission), showed large interspecies variation, ranging from <310nm (European mole) to 494nm (European ground squirrel) (median 401.5 nm). It is well established that age can alter lens coloration and size, thereby modifying filtering properties(29). Thus, human pre-receptoral filtering standards are corrected for age(21). Given the challenges associated with accessing eyes of different ages from different species, we opted to perform our calculations using adult lens filtering for each species.

### 2.2. Calculation of species- and photoreceptor-specific light exposure metrics

Having determined methods for estimating *in vivo* spectral sensitivity we applied them to define functions for calculating α-opic irradiance for 12 domestic mammal species. The method of quantifying α-opic irradiance is captured by **equation 1** (**Table 2**) and relies on a full description of s_α,s_(λ), the *in vivo* spectral sensitivity of the target photoreceptor (α) in target species (s). We defined s_α,s_(λ) for rod, cone and melanopsin photoreceptors across the 12 species of domesticated mammals by using the opsin template of Govardovskii and colleagues(30) and λ_max_ values from **Table 1** to produce a full *in vitro* spectral efficiency profile. We then multiplied by lens transmittance to provide s_α,s_(λ), our estimate of *in vivo* spectral sensitivity for each pigment. Full functions for all pigments in all species are available in **Supplementary data 2A**.

**Table 2:** Formulas used to calculate species and photopigment-specific light exposure metrics.

|  |  |  |
| --- | --- | --- |
| 1 | Species-specific α-opic irradiance | $E_{e,\alpha,s} = \int E_{e,\lambda}(\lambda) s_{\alpha,s}(\lambda) d\lambda$ |
| 2 | Species-specific α-opic efficacy of luminous radiation | $K_{\alpha,s,v}^{D65} = \frac{E_{e,\alpha,s}^{D65}}{E_v^{D65}}$ |
| 3 | Species-specific α-opic equivalent daylight illuminance | $E_{v,\alpha,s}^{D65} = \frac{E_{e,\alpha,s}}{K_{\alpha,s,v}^{D65}}$ |

Descriptions of α-opic irradiance for humans increasingly employ a derived quantity termed α-opic equivalent daylight illuminance (EDI)(31). EDI represents the illuminance (units = lux) of a standard daylight spectrum (termed D65) that would provide the equivalent α-opic irradiance. The method for calculating species-specific α-opic EDI (s α-opic EDI; e.g. ‘mouse melanopic EDI’), involves first determining α-opic efficiency of D65 (K^D65^ ; W/lm) (**Table 2, Formula 2**) and then dividing α-opic irradiance (E_e,α,s_) by this value (**Table 2, Formula 3**). For simplicity, K^D65^ is provided for all target photoreceptors in **Supplementary data-2B**.

We provide two resources to facilitate calculation of species specific α-opic irradiance and EDI according to the equations in **Table 2** and the s_α,s_(λ) functions in **Supplementary data 2**: an R package (alphaopics), which includes functions for calculating species and opsin-specific units (https://doi.org/10.48420/23283059); and an online toolbox (Alphaopics: Species-specific light exposure calculator) for easy calculation of species-specific metrics (https://alphaopics.shinyapps.io/animal_light_toolbox/). Both require the user to provide a measure of spectral power distribution (E_e,λ_(λ) for the light reaching the animal’s eye.

### 2.3. Architecture for a miniaturised mammalian α-opic meter

Calculating α-opic quantities from full spectral power distributions requires an understanding of the properties of light and investment in appropriate measurement technology. For most users, a more ‘point and click’ solution to light measurement is required. Widely available lux meters already provide this when measuring photopic illuminance. Lux meters generally achieve the appropriate spectral weighting by combining the spectral sensitivity of the light sensor with an optic filter with an appropriate spectral transmission. Sadly, that strategy is not scalable for the α-opic metrology because, in principle, separate filters may be required for each potential target photoreceptor in each species. We wondered whether multichannel miniaturised (MM) light sensors could provide a solution to this problem and form the basis of easy-to-use light meters recording species specific α-opic units. MM sensors comprise 6 or more detectors, each sitting below an independent narrowband optical filter. They are relatively cheap, have high measuring accuracy of around 90% against calibrated sources, and have been used successfully to estimate human α-opic metrics(32–34). We set out to determine whether this technology could form the basis of accessible species-specific light meters. To this end, we attempted to recalibrate an open-source wearable light dosimeter (SpectraWear)(35) based upon a 10 channel MM sensor chip (AMS AS7341, Premstaetten, Austria) for species specific measurements.

To facilitate estimation of species- and photopigment-specific light exposures, we generated a set of narrow- and broad-band light stimuli with energy spanning the visible range (**Figure 2A**; see Methods). We then measured these with a spectroradiometer and applied the ‘alphaopics’ package to calculate species-specific α-opic EDIs. Next, we used nonlinear least-square fitting to derive weighting coefficients for the 10-channel sensor readings from SpectraWear that best recreated the species-specific α-opic EDIs provided by those stimuli. We finally validated the resulting calibration coefficients against a test set of lights generated from the narrow- and broad-band sources used for calibration but spanning a wider range of irradiances. A comparison of measured (based upon full spectral power density measurements) and predicted (based upon SpectraWear) EDIs showed strong correlations for all α-opic irradiances in a single representative species (**Figure 2B**). To provide a more comprehensive description of the device performance across α-opic quantities and species, we calculated the estimation error (difference between measured and predicted α-opic EDI) for each test stimulus for the α-opic quantities of our 12 domestic mammal species and humans (**Figure 2C)**. This revealed variations in performance across different α-opic quantities, with consistently high accuracy for melanopic and rhodopic EDIs for all species evaluated (**Figure 2C**; typical estimation errors = 0.06±0.01 & 0.05±0.01 log units respectively; median±SD). Despite the substantial variation in cone opsin λ_max_ across species, L- and M-cone opic EDIs were also estimated with good reliability (**Figure 2D**; typical estimation errors = 0.06±0.02 log units). Conversely, performance was reduced for S-cone-opic EDIs, especially in species where S-cones show peak sensitivity in the UV (**Figure 2D**; typical estimation errors = 0.30±0.45 log units). In sum, the performance of the device allowed us to reconstruct reliable estimates (within ±17%) of α-opic EDIs other than S-cone opic. This technology could enable continuous light monitoring in field with a scalable design.

**Figure 2:**
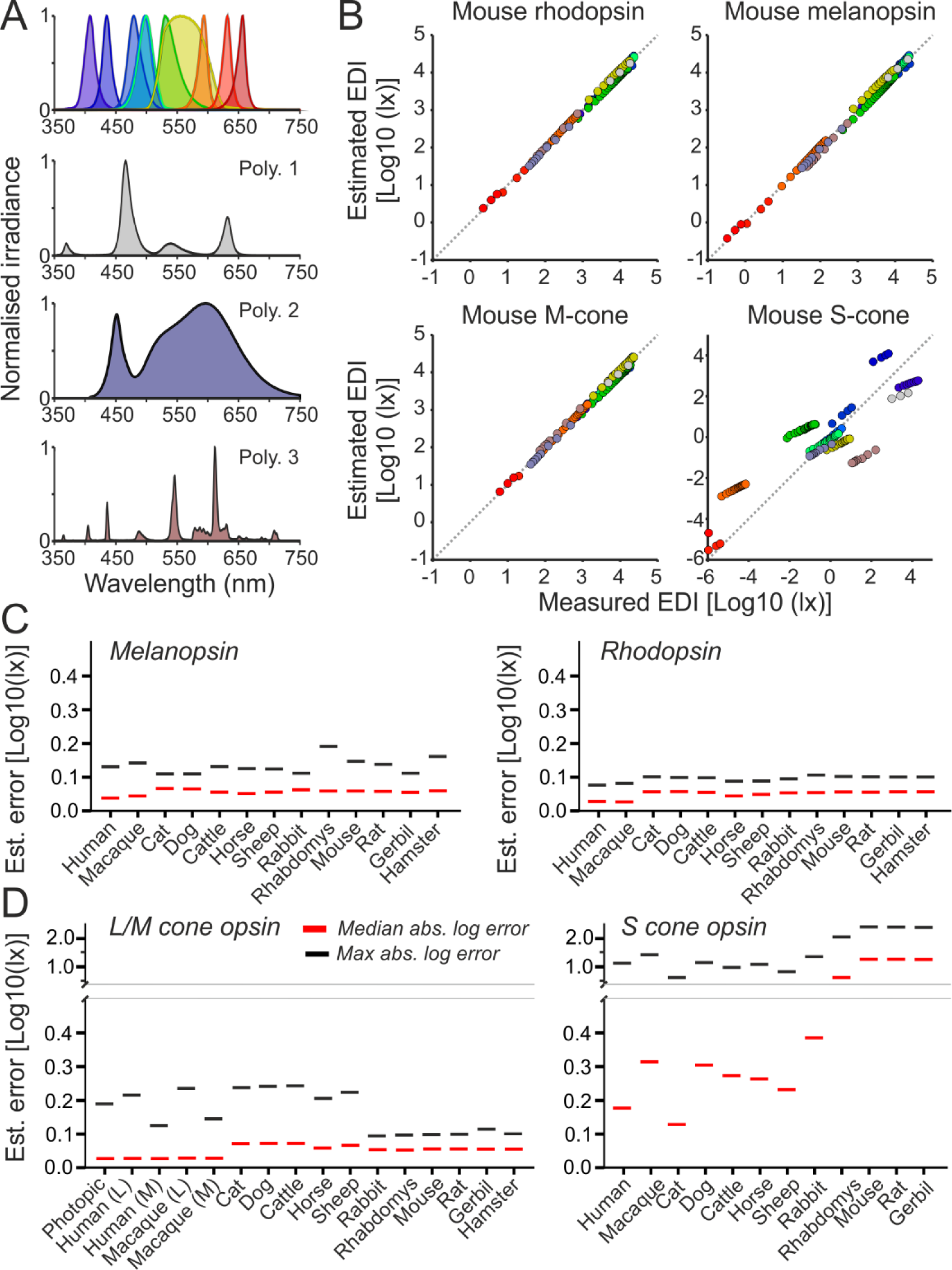
Cross-species light dosimeter validation. (**A**) Normalised spectral power distributions of ten narrow-(top) and three broadband stimuli (lower panels) used for device calibration and validation. (**B**) Top panels show scatter plots of relationship between mouse α-opic EDIs, determined based on spectroradiometric measurements, for stimuli in A across a range of irradiances and the corresponding estimated melanopic EDIS based on weighted readings from the 10-channel light sensor. (**C,D**) Plots showing median and maximum log absolute errors for melanopic (**C**, left), rhodopic (**C**: right), L/M-cone opic (**D**, left) and S-cone opic (**D**, right) EDIs across species.

### 2.4. Characterisation of species-specific light exposure amongst common illuminants

We finally turned to describing the suitability of the α-opic metrology for predicting the response of animals to light of divergent spectral composition. In the first instance we asked whether α-opic units predicted responses within a single candidate species to spectrally divergent stimuli. That is a prerequisite for using any metric to standardise husbandry or recreate experimental conditions. Many studies of circadian photoentrainment in mice quantify light in photopic lux. We took four datasets describing irradiance response curves for circadian phase shifting in wild-type and retinal degenerate mice across the wavelength range(36–38) and expressed them either as a function of photopic lux, or the mouse α-opic EDIs (**Figure 3A, B, Supplementary figure 2A-C**). We found that the fraction of variance in circadian phase shift predicted by light intensity (R^2^ for curve fit) was >0.8 when light intensity was quantified in any of mouse melanopic, rhodopic or M-cone-opic EDIs but substantially reduced when either photopic lux (0.4) or S-cone opic EDI (0.2) were used (**Figure 3C**). This finding highlights the superior capacity of the α-opic metrology to predict mouse circadian phase resetting to spectrally divergent lights.

**Figure 3:**
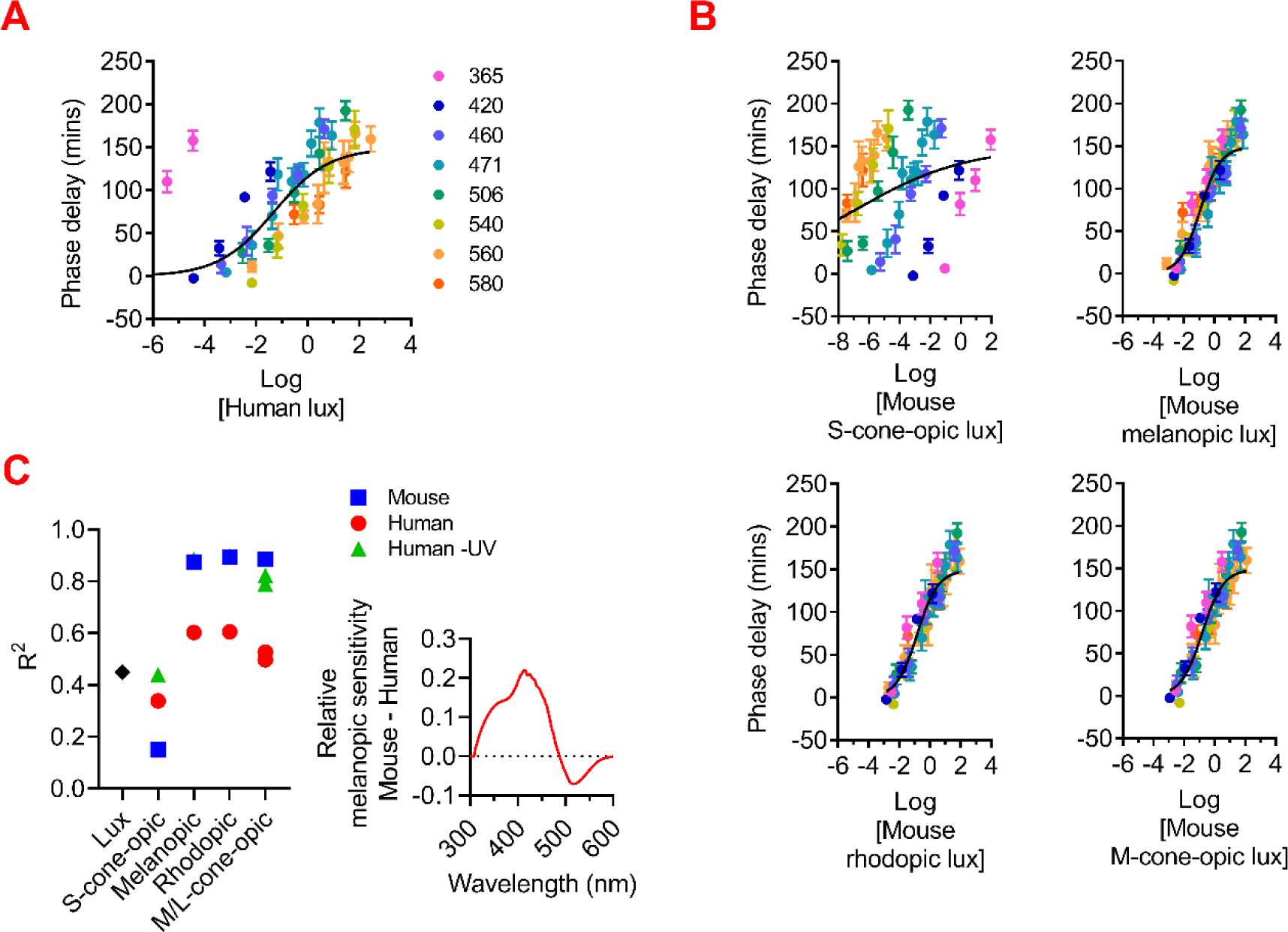
Irradiance response curves for circadian phase shifting in C57 wild-type mouse(37). Phase delays (mean±SEM) were plotted against eight narrowband light stimuli with a range of intensities. Light stimuli were presented as human photopic lux in (**A**), mouse α-opic EDIs in (**B**). Non-linear four-parameter fit lines were shown in all plots. R^2^ for curve fits were shown in (**C**). In addition to mouse-specific α-opic EDIs, curve fits for human α-opic EDIs were also presented (with and without light stimuli at 365nm). Lower right plot shows comparison of relative melanopic sensitivity for mouse vs human as a function of wavelength.

We wondered the extent to which this improvement relied upon adoption of species-specific metrics or whether human α-opic units would achieve the same effect. To this end, we generated versions of the mouse irradiance response curves with light intensity expressed in human α-opic units. As there is great cross-species divergence in spectral sensitivity of cone photoreceptors, it is perhaps unsurprising that human L/M-cone-opic EDIs provided an inadequate prediction of the mouse response (**Figure 3C**). The reduction in goodness of fit when expressing light in human melanopic or rhodopic EDI is less expected given the similarity in spectral sensitivity of melanopsin and rod opsin photopigments across mammals. However, there is substantial divergence in lens transmission to short wavelength light across these species. Accordingly, an assessment of relative melanopic sensitivity for mouse vs human as a function of wavelength (**Figure 3C**) reveals that, while the two species have very similar sensitivity >450nm, their sensitivity to shorter wavelengths diverges. It follows that human quantities may be more appropriate for lights that lack strong output at very short wavelengths (as is the case for most artificial sources). We tested this prediction by excluding data for UV wavelengths from our phase shift dataset and recalculating goodness of fit to human melanopic and rhodopic EDI. In both cases the human metrics now provided goodness of fit for the mouse phase shifting irradiance response curve (**Figure 3C**). Together these analyses reveal the advantages of using species-specific versions of the α-opic units, while showing that there may be circumstances (melanopic and rhodopic EDIs for stimuli with little UV output) under which human metrics are an acceptable alternative.

Having confirmed the advantages of the α-opic metrology for predicting the animal response to light, we quantified the potential error associated with the current practice of quantifying light in photopic lux. An appropriate measurement system should allow the animal’s experience of lights differing in spectral composition to be normalised. We therefore took α-opic EDI as a measure of true effective intensity and asked how well the current practice of using measuring light in photopic lux predicted α-opic EDI. We started with the most widely encountered case of broad-spectrum lights of the types used for general illumination applications. Taking 42 such broad-spectrum lights, intensity matched for photopic lux (and thus under current practice considered to be interchangeable for animal work), we calculated their α-opic EDI for each of the 13 mammals. This revealed substantial variation in effective intensity for each photoreceptor (α-opic EDIs) across the various light sources. Shown for a representative species (mouse) in **Figure 4A** left. At the extreme, two broad-spectrum lights, intensity matched in photopic lux, could show 85% difference in melanopic EDI. The poor suitability of the photopic lux metric for predicting α-opic EDI was even starker for monochromatic or ‘coloured’ lights (**Figure 4A** right).

**Figure 4:**
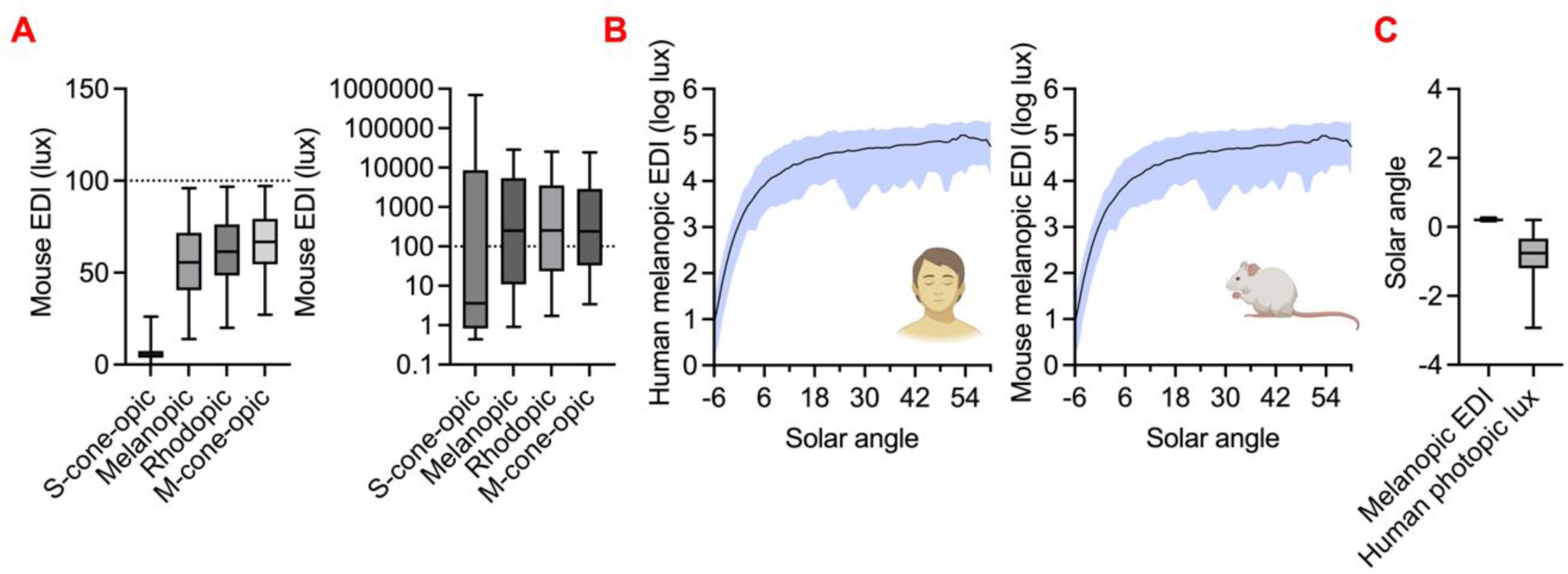
Cross-species α-opic EDIs across different light sources. (**A**) Comparison of mouse α-opic EDIs across (left, linear Y-axis) 42 broad-spectrum CIE standard white light sources and (right, log-scale Y-axis) 9 monochromatic LED light sources matched for 100 human photopic lux. Box plots show mean and ranges. (**B**) Plots of the relationship between solar angle and human (left) and mouse (right) melanopic EDIs for a range of real-world light measures. Natural spectral irradiances over 16 days were collected in the Netherlands (latitude: 53.24°, longitude: 6.54°, summer daylengths >15h) and comprises overcast and clear weather conditions shown with error range(60). (**C**) White light sources in **A** were converted to species-specific melanopic EDIs for 13 species reported in this study. Box plots show mean±range of solar angle that represent either 1000 species-specific EDI lux or 1000 human photopic lux matched light inputs.

Thus far, we have considered the advantages of the α-opic EDI metrology for quantifying effective intensity for different light sources within a species. A further aspect of the metrology is its ability to allow informative comparisons of light intensity between species, e.g., to ask whether some species are fundamentally more sensitive to light than others. The α-opic EDI concept incorporates an anchor to the natural world (an equivalent amount of daylight), which should facilitate ecologically relevant comparisons cross-species. To illustrate this, **Figure 4B** shows a plot of the relationship between solar angle and human melanopic irradiance for a range of real-world light measures. The nature of the correction used to convert α-opic irradiance to EDI ensures that this relationship is near identical for all α-opic EDIs across all species (r>0.99). It follows that lights of equivalent α-opic EDI should recreate each animal’s experience of a similar solar angle with good accuracy even across species. To confirm this, we calculated the solar angle corresponding to 1000 lux melanopic EDI in each of 13 mammalian species (humans + the 12 domesticated mammals defined above). The result is plotted in **Figure 4C** and confirms that in all species this recreates the experience of natural light when the sun is just above the horizon. For comparison we estimated the appropriate solar angle for our 42 broad-spectrum lights when set to 1000 photopic lux (by converting to melanopic EDI and relating to the function in **Figure 4B**). In this case, the equivalent solar angle became more variable, encompassing sunrise and much of the civil twilight range (**Figure 4C).**

## 3. Discussion

No animal is equally sensitivity to light at all wavelengths. It follows that applying the appropriate spectral weighting function is critical in quantifying light in photobiology. The α-opic concept of light measurement was first proposed in 2014 to account for the then still quite recent discovery of the inner retinal photoreceptor melanopsin, and the related realisation that lights matched for photopic illuminance could have quite divergent ability to elicit important circadian and neurophysiology light responses(20). The α-opic metrology has a quite different conceptual basis than conventional photometry. Whereas photopic illuminance quantifies light according to the spectral sensitivity of a single distinct percept (perceived brightness under cone favouring conditions), the α-opic metrology aims to quantify light according to the experience of each photoreceptor type, remaining agnostic to its final application in supporting vision or reflex light responses. What is lost in simplicity is gained in flexibility. Thus, while the single measure of photopic lux is replaced by 5 α-opic irradiances (for humans), the latter capture all relevant information about incident light, whereas the former only captures information about its perceived brightness. The additional flexibility of the α-opic metrology is especially valuable when quantifying light for non-human animals. Most mammals lack the long-wavelength shifted L/M cones that dominate the photopic sensitivity function in humans(19). Replacement spectral efficiency functions for perceived brightness are not readily obtainable for many species and in any case would not capture important circadian and neurophysiological light responses. By contrast, information about photoreceptor spectral sensitivity is often available, allowing species specific α-opic functions to provide a holistic description of the animal’s experience of light. This ability of the α-opic metrology to provide species-specific quantification of light was recognised in the initial description of the approach and yet, while α-opic units are now increasingly presented in the context of human light exposure(31) they have not been widely adopted for non-human animals. There are likely several reasons for this. Most importantly, a push for more widespread appreciation of this strategy has awaited formal specification of the α-opic approach as an internationally approved standard metrology. With that in place, more practical problems about how to measure species-specific α-opic units come into focus. Here we have addressed two of these: gaps in our knowledge of photoreceptor spectral sensitivity for some species; and a high barrier to entry for those wishing to measure light in this way. Having addressed these, we provide some simulations to quantify the advantages of applying α-opic units in animal biology.

Our approach to filling gaps in knowledge of photoreceptor spectral sensitivity has been to employ a heterologous action spectroscopy approach, which we first used to study human melanopsin(23). We have adapted that strategy to make it higher throughput by optimising the range of wavelengths/irradiances used and by applying a bootstrap modelling approach to analysis which provides the optimal pigment λ_max_ for the data collected and an error estimate for that value. The result is a relatively high-throughput method, requiring little specialist lab equipment and applicable to any photoreceptor with known cDNA sequence. We have applied it to define melanopsin spectral sensitivity because, while other methods (electroretinography or microspectrophotometry) are available for rods and cones in domestic mammals, studying melanopsin *in vivo* is more challenging(27, 39). In principle the method could be applicable to any photoreceptor from any species for which a live cell readout of photoactivation is available. One important consideration in these data is that cDNA sequences for melanopsin are not available for many species. In these instances, predicted coding sequences for genomic data have been used, with the associated potential for errors(40, 41). Indeed, when aligning melanopsin sequences against the confirmed sequence for humans, several sequences listed on databases were missing key regions due to mis-identified splicing. It is for this reason that we did not rely upon one gene database for the generation of melanopsin sequences.

Our data reveal conservation of melanopsin spectral sensitivity across the 13 mammalian species described here. The total range of predicted λ_max_ is 15nm, which is similar to that for rod opsin across these species (13nm), but small compared to that of either S- or M-cones (>50nm). The fact that the species showing greatest difference in melanopsin λ_max_ are both small diurnal mammals from semi-arid environments (striped mouse and mongolian gerbil) adds to the impression that this parameter is not under divergent selection pressure across the species studied here. An interesting question is what features of the light environment and/or structural contraints for the protein may be responsible for restricting melanopsin λ_max_ to ∼480nm.

Opsin photopigments can employ a range of cis-isoforms of retinaldehyde as chromophore, and in the case of melanopsin there is evidence of diversity in the choice of isoform *in vivo*. As opsin spectral sensitivity is influenced by the retinaldehyde isoform used, it is important to note that the values presented here represent those with the 11-cis isoform of A1 retinaldehyde. The resultant estimates for spectral sensitivity match those for whole animal responses in mice, macaque and humans. The reader is directed elsewhere for a complete consideration of factors determining estimates of melanopsin spectral sensitivity in other elements of the literature(42).

In principle, the method of calculating α-opic EDI is applicable to any photoreceptor from any species. In the case of mammals, future work may extend it to Opn5 and/or Opn3 as further evidence of their sensory functions accumulates, although careful consideration of appropriate pre-receptoral filtering is required for opsins expressed outside of the eye(43–45). The number of α-opic EDIs required to fully quantify the light environment for many non-mammalian species may be large, as these commonly have many photopigment types expressed in different parts of the body (and thus subject to divergent pre-receptoral filtering). Nevertheless, calculating these quantities would represent an advance on alternatives that either assume equal sensitivity across the spectrum (unweighted sum of energy/quanta) or human spectral sensitivity (photopic illuminance).

Spectrometers capable of providing spectral power density measures, which in combination with a suitable wavelength weighting function can be used to calculate species-specific α-opic metrics, are widely available. The toolbox presented here to facilitate this process represents an extension on previously published versions restricted to a smaller number of species(17, 20, 46, 47). More accessible ‘point and click’ solutions to calculate species specific units could take advantage of meters recently developed to measure human α-opic metrics. In particular, we show here that appropriate calibration allows the MM technology forming the basis of several such meters to measure many species specific quantities with acceptable accuracy (<17% error rate). We validate an approach based upon a system developed an open hardware principles (Spectrawear(35)), but commercially available products could in principle be adapted to this purpose. One important caveat here is that core MM chips generally do not have good coverage at short wavelengths (especially UV) over which many domestic mammals are much more sensitive than humans(18, 48). That likely explains the poor performance of Spectrawear for S-cone opic EDI and suggests that meters including separate UV sensitive detector(s) could have superior performance.

Quantifying light in species-specific α-opic EDI has clear conceptual advantages over current practices of using either photopic illuminance or total energy/quanta. We show here that it also provides superior ability to predict circadian phase shift responses in mice (across numerous studies) and allows sensible comparisons of effective light intensity across species. Application of this metrology could thus bring coherence to the growing literature on light effects on mammalian physiology and behaviour and reproducibility to any experiment in which light influences the outcome. Appropriate measurement can also have a wider significance for animal welfare. Insufficient daytime light and excessive light at night have been shown to disrupt circadian rhythm and sleep, and have negative impacts on the health of animals, as well as research outputs and scientific reproducibility(4–12). Animal-centered approaches are key to enhancing the health and wellbeing of indoor housed animals, and the accurate provision of light is a crucial consideration. Environmental light pollution poses a significant disruptor to many animal ecosystems, emphasizing the need for better characterization of animal-specific light exposure to improve conservation strategies(49, 50). Furthermore, given the significant land and energy usage required for farm animal operations, the identification of optimum lighting conditions that balance productivity, health, and electricity usage has the potential to generate substantial energy savings(51–53). Additionally, the impact of evening and night-time light exposure in the home environment on human sleep is well-documented(54), but remains unknown for pets(55). One process that could facilitate these applications would be simplification of the 4 α-opic quantities required to fully describe irradiance for most mammalian species to a single metric that provides a reasonable prediction of light responses of interest under most circumstances. This could be a single α-opic metric or a composite of several. Such a process has led to the increasing use of melanopic EDI as a single metric for non-visual light responses in humans(31).

## 4. Conclusions

Our study reveals an accessible method to measure photopigment-specific “α-opic” light exposure for mammal species. We present the prerequisite data for defining α-opic metrics; lens transmission, and novel action spectra for melanopsins from most major domesticated mammalian species. We then present the necessary calculations to derive photoreceptor-specific metrics and provide open access software for easy calculation. Our data reveals that species-specific α-opic metrics offer greater accuracy for the description of the physiological effects of light than the current commonly used standard of photopic lux. Finally, we present a prototype low-cost and scalable portable light dosimeter for the measurement of lighting conditions. This method for light measurement allows for the easy monitoring, regulation and intervention of light exposure in animal housings and will lead to increased research accuracy using animal models, agricultural efficiency and improve animal health and wellbeing.

## 5. Methods

### 5.1. Recombinant cloning of animal opsins

Coding sequences for mammalian melanopsins were accessed from either NCBI GenBank or Ensembl databases (Ensembl Release 109(56)). Open reading frames for the following sequences were used to construct expression vectors: brown rat Opn4, NM_138860.1; cat Opn4, NM_001009325.2; cattle Opn4, NM_001192399.1; crab-eating macaque Opn4, ENSMFAT00000002526.2; dog Opn4, XM_038662366.1; four-striped grass mouse Opn4, in house cDNA; horse Opn4, XM_023648726.1; human OPN4, NM_033282.4; Mongolian gerbil Opn4, XM_021635996.1; mouse Opn4L, NM_013887.2; Rabbit Opn4, ENSOCUT00000017574; Sheep Opn4, XM_027962232.2; Syrian hamster Opn4, ENSMAUT00000015782 (**Supplementary table 3, 4**). Gene sequences were synthesised using ThermoFisher GeneArt Gene Fragment synthesis and TwistBio Gene Fragment synthesis and underwent codon optimisation where necessary for synthesis. All opsin sequences were tagged with the 1D4 epitope (TETSQVAPA) on the C-terminus. Opsins were introduced into the multiple cloning site of the pcDNA3 vector (Invitrogen) downstream of the CMV promoter using NEBuilder HiFi Assembly (New England Biolabs).

### 5.2. Heterologous action spectroscopy

HEK293T cells (American Type Culture Collection) were cultured in Dulbecco’s modified Eagle’s Medium (4.5 g l−1 D-glucose, sodium pyruvate and L-glutamine with 10% foetal calf serum; DMEM). Cells were transiently transfected with 500ng plasmid expression vectors for the relevant opsin and 500ng genetically encoded Ca^2+^ indicator mtAequorin (as described in (23)) using lipofectamine 2000 (Invitrogen) and incubated overnight with 10µM 9-*cis*-retinal (Sigma-Aldrich) or 10uM 11-*cis*-retinal (National Eye Institute, National Institutes of Health). The following day, cells were incubated with 10µM Coelenterazine-h (Promega) in the dark for 2 hours before recording luminescence in a plate reader (Optima FLUOStar, BMG) modified to allow “In-well” stimulation with an external light source (CoolLED pe-4000, CoolLED) via fibre optic. Luminescence recordings were sampled at a temporal resolution of 2 seconds per timepoint. Baseline luminescence was recorded for 10 seconds, after which cells were stimulated with light (1s duration) of varying intensities (11 – 16 log photon/cm^2^/sec total photon flux) at one of 6 different wavelengths (435nm, 460nm, 470nm, 490nm, 500nm, 525nm).

### 5.3. Calculation of opsin photon sensitivity peaks

To determine the λ_max_ values for each opsin, we employed a nonlinear optimization strategy. In this strategy, an optimization algorithm systematically iterates over different values of λ_max_ (*optim* function in *R* (version 4.3.0), using the Brent search method). Each iteration consisted of two steps. First, the effective photon flux values of the light sources were updated according to the Govardovskii photopigment template(30) corresponding to the currently assumed value for λ_max_. Then, a 5-parameter log-logistic model was fitted with cell response and the updated effective photon flux as dependent and independent variables respectively (*drm* function from the *drc* package (version 3.0-1)), from which the estimation error was extracted (i.e., residual sum of squares). The optimization algorithm searched for the λ_max_ value (within a 400-600 nm range) that would minimize this error. Finally, bootstrapping was performed in which the above optimization procedure was repeated 1000 times, each time using only a random subset of the data (with replacement). The average and standard deviation of the 1000 resulting λ_max_ values were finally used as the λ_max_ estimate and estimation error.

### 5.4. Standardization of animal lens transmissions

Literature searches for lens, cornea, and vitreous humor light transmissions for mammal species were performed. We accessed data from 56 adult species. References and data are listed in **Supplementary data 1**. Human pre-receptoral filtering is based on a reference observer of age 32 years(20, 21). If available in the reference source table or supplement, original data was used. Otherwise, data was extracted from reference plot using WebPlotDigitizer (Version 4.6). If relative absorbance was reported, transmission was calculated as 100 × 10^−*Relative absorbance*^. To harmonize the data the following steps were performed. If less than 50 data points available, cubic splines with 1 nm step were interpolated. If more than 50 data points available, smoothing splines with 1 nm step and 50 knots were interpolated. All raw data were normalized to their maximum values (max. 100%). Data were filled with the last value until 800 nm and were filled until 310 nm using the slope of the first 3 values. Negative values were accepted as zero. Where multiple valid sources were available (e.g., Syrian hamster and Brown rat), mean fits were used. For seven species (European ground squirrel, Syrian hamster, thirteen-lined ground squirrel, coruro, Mongolian gerbil, seal, cattle) cornea or vitreous humor filtering information were available. For those, we compared the wavelength where transmission reached 50% in lens, cornea, and vitreous humor. Lens transmission for European ground squirrel (Spermophilus citellus) and Syrian hamster were measured as previously described(57), for tree shrew (Tupaia belangeri) and Fat-Tailed Dunnart (Sminthopsis crassicaudata) an AvaSpec 2048 (Avantes) UV/VIS spectrometer with a perpendicular fibreoptics transmission setup was used.

### 5.5. Prototyping of a mammal light dosimeter and its calibration

The prototype is based on an open-access human α-opic light dosimeter electronic design(35). The device prototype of 3D-printed black plastic outer case, micro SD card memory storage and Bluetooth control. The device light detection had transparent acrylic disc (Perspex) with 20mm diameter and with a diffuser (Optsaver L-35 Kimoto, Cedartown, Georgia, USA) underneath. The device incorporates AMS AS7341 multichannel spectral colour sensor (ams, Premstaetten, Austria), which had channels having peak wavelengths at 415nm, 445nm, 480nm, 515nm, 555nm, 590nm, 630nm, 680nm, 910nm, and a clear channel to read unfiltered spectral input. Sensor reading of a prototype device were collected across 169 light conditions (13 distinct spectra across multiple irradiances). This included ten distinct narrowband spectra, generated via a calibrated multispectral LED light source (CoolLED pE-4000 LED Illumination System; narrowband peaks: 405, 435, 460, 470, 490, 500, 525, 550, 595, 635 and 660nm) and three distinct broadband spectra (Philips CorePro white LED 470 lumen 4000K, Philips Tornado white fluorescent 1570 lumen 2700K or CoolLED pE-4000 LED Illumination System 365-460-525-635nm colour-mixed white LED) Throughout, stimuli were measured via a calibrated spectroradiometer (SpectroCal, Cambridge Research Systems, UK) and converted to species specific α-opic EDIs as described above. All measurements were performed in a dark room. We then collected an identical set of measurements of the same stimuli user our 10-channel light sensor (integration time 182ms, automatic gain optimisation in the range of 8 – 512X and additional post hoc scaling by a factor of 10^6^ such that sensor counts took on positive values >=1). To calibrate the device, we then selected a subset of 3 measurements for each of the 13 distinct spectra described above. Based on the known relationship between the measured sensor counts and the α-opic irradiance of these calibration stimuli (and our previous observations that these sensors exhibit good linearity across a very wide range(35)), we extrapolated the expected sensor counts for each spectra across a consistent set of EDI values (-1, 0.5 and 2 log lux). We then fit a set of weighting coefficients such that the sum of the adjusted log sensor counts best recreated the expected log EDIs across stimuli in the calibration dataset (using ‘lsqcurvefit’ function in Matlab R2018a, Mathworks, MA, USA). We choose to perform fits on log transformed data since this allowed for sensor weightings to be either positive or negative (important for reliable estimates from low channel count sensors such as this) while avoiding the possibility that any resulting estimated α-opic EDI might take on (impossible) negative values. To validate the resulting species- and photoreceptor-specific device calibration, we then used the derived sensor weighting coefficients to estimate EDIs across the remaining 130 spectroradiometrically measured test stimuli that didn’t contribute to calibration (n=3-27 irradiances/spectrum at unweighted irradiances of ∼0.2-80W/m^2^). Log (absolute) errors for these estimates, relative to the directly measured values, were then determined for each distinct spectrum, species and photopigment.

### 5.6. Characterisation of species-specific light exposure amongst common illuminants

We selected four datasets(36–38) including more than five light stimuli with different spectral distributions. Phase shift values were extracted from graphs using WebPlotDigitizer (Version 4.6) and light stimuli spectral power distributions were generated as normal distribution with specified peak wavelength and half width at half maximum values (Matlab R2018a, Mathworks, MA, USA). We then converted them to human photopic lux, the mouse α-opic EDIs and the human α-opic EDIs. We fitted non-linear four-parameter lines to estimate phase shifts using light stimuli.

All test light sources were arbitrarily set to 100 human photopic lux. Indoor artificial standard illuminants included: CIE standard illuminant A (incandescent 2855 K), CIE standard illuminant HP types (High pressure sodium lamps 1-5; standard, colour-enhanced, metal halide), CIE standard illuminant FL types (Fluorescent 1-12 and 3.1-3.15; standard, broad-band, narrow-band, standard halophosphate, DeLuxe type, three-band, multi-band, D65 simulator), CIE standard illuminant LED types (Light-emitting diode B1-B5, BH1, RGB1 and V1-V2; Phosphor-type LEDs with different correlated colour temperatures, Hybrid-type, RGB-type, and violet-pumped phosphor-types)(58, 59). As narrowband test light source, we measured spectral power distributions of CoolLED pE-4000 LED Illumination System (narrowband peaks: 365, 405, 435, 470, 500, 525, 550, 595, 635) using a spectroradiometer (SpectroCal, Cambridge Research Systems, UK). For 13 species which we have both opsin sensitivity and lens transmission information (Human, mouse, four-striped grass mouse, brown rat, Syrian hamster, Mongolian gerbil, cattle, sheep, horse, cat, dog, crab-eating macaque, rabbit), α-opic EDIs were calculated. Using GraphPad Prism 9, between-species mean and range were plotted for each light source.

Natural daylight spectral irradiances, solar angle (degree) and weather conditions on multiple days were collected in the University of Groningen, the Netherlands (latitude: 53.24°, longitude: 6.54°)(60). Daylengths at the measurement date were calculated using R package ‘suncalc’ (0.5.1). We used a subset of the data (−6° to 60° solar angle, weather conditions >6 cloudy or <3 clear, summer daylengths >15h). In total, our data included 5 days of clear sunny conditions (4633 measurements) and 11 days of overcast daylight (10433 measurements). Human and mouse melanopic EDIs (mean ± range) were plotted against solar angle. For 13 species, α-opic EDIs representing that solar angle were calculated and compared pairwise using Pearson correlation. Finally, above mentioned CIE light sources matched for either 1000 human photopic lux or species-specific melanopic EDI lux (for 13 species). These values were converted to solar angles using the above mentioned curves.

## 6. List of Abbreviations

9-*cis*: – 9-*cis* retinaldehyde
11-*cis*: – 11-*cis* retinaldehyde
λ_max_: – The wavelength at which maximum activity/absorption occurs
EDI: - equivalent daylight illumination
MM light sensor: - multichannel miniaturised light sensor
UV: - Ultraviolet

## 7. Declarations

### Ethics approval and consent to participate

Transmission data was collected under the University of Groningen Animal Experiments Committee license number BG02197/98.

### Consent for publication

Not applicable.

### Availability of data and materials

The datasets generated and/or analysed during the current study are available in the following Figshare repository: https://doi.org/10.48420/23708610. Functions for calculating the Functions for calculating species and opsin-specific units are available as an R package (alphaopics) (https://doi.org/10.48420/23283059); and an online toolbox (Alphaopics: Species-specific light exposure calculator) for easy calculation of species-specific metrics (https://alphaopics.shinyapps.io/animal_light_toolbox/).

### Author’s contributions

RJM and RJL designed the opsin sensitivity measurement methodology, and RJM and MJG performed the photopigment sensitivity experimental measurements. AD searched literature for prereceptoral filtering data. RJM searched literature for spectral sensitivities. RAH performed novel transmission measurements. AD, RJM, TW and TMB performed data analysis. TMB, RJL and AD were involved in the design, production and calibration of the light dosimeters. AD, TW, RJM and RJL drafted the manuscript. TMB, RJL, AD and RJM were involved in planning and supervising the project. All authors discussed the results and commented on the manuscript. All authors have read and agreed to the final manuscript.

## Acknowledgements

We thank Lucien Bickerstaff for his help in calibration of the light dosimeter. We also thank Prof Ron Douglas (City University London) for his guidance on lens transmission of mammals.

## Additional Files

Supplementary Table 1: Literature search for opsin spectral sensitivities in the absence of prereceptoral filtering.

Supplementary table 2: Different isoforms of retinal in measurements of OPN4 spectral sensitivities (λmax).

Supplementary table 3: Amino acid sequences of mammalian melanopsins used in study.

Supplementary table 4: Nucleotide sequences of mammalian melanopsins used in study.

Supplementary data 1: Literature search for lens transmission for 56 mammalian species.

Supplementary data 2A: Estimation of *in vivo* spectral sensitivity for each photopigment for each mammalian species.

Supplementary data 2B: α-opic efficiency of D65 (K^D65^ ; W/lm) for each photopigment for all target species.

